# Analysis of the intrinsic chromatin binding property of HIV-1 integrase and its regulation by LEDGF/p75 using human chromosomes spreads

**DOI:** 10.1101/2020.12.23.424107

**Authors:** D. Lapaillerie, E. Mauro, B. Lelandais, G. Ferran, N. Kuschner, C. Calmels, M. Métifiot, C. Rooryck, M. Ruff, C. Zimmer, P. Lesbats, J. Toutain, V. Parissi

**Author notes:** International Associated Laboratory (LIA) of Microbiology and Immunology, CNRS / University de Bordeaux / Heinrich Pette Institute-Leibniz. Viral DNA Integration and Chromatin Dynamics Network (DyNAVir).

## Abstract

Retroviral integration requires the stable insertion of the viral genome into the host chromosomes. During this process, the functional integration complex must associate with cellular chromatin *via* the interaction between retroviral integrase and nucleosomes. The final association between the HIV-1 integration complex and the nucleosomal target DNA remains unclear and may involve the chromatin-binding properties of both the retroviral integrase and its cellular cofactor LEDGF/p75. To date, there is no experimental system allowing the direct monitoring of this protein association with chromatin to depict the molecular mechanism of this process fully. To investigate this and understand the LEDGF/p75-mediated chromatin tethering of HIV-1 integrase further, we used both biochemical approaches and an unedited chromosome-binding assays. Our study revealed that retroviral IN has an intrinsic ability to bind and recognize specific chromatin regions even in the absence of its cofactor. We also showed that this integrase chromatin-binding property was modulated by the interaction with its cofactor LEDGF/p75, which redirected the enzyme to alternative chromatin regions. Using these approaches, we also better determined the chronology of efficient LEDGF/p75-mediated targeting of HIV-1 integrase to chromatin. In addition to supporting a chromatin-binding function of the integrase protein acting in concert with LEDGF/p75 for the optimal association with the nucleosomal substrate, our work precisely elucidates the mechanism of action of LEDGF/p75 in this crucial integration step.

## INTRODUCTION

Retroviral infection requires the stable insertion of the viral DNA genome into the chromosomes of the host (for a review on integration, see (Lesbats et al. 2016)). This interaction between the incoming integration complex and the host chromosome thus constitutes the first contact between the viral and cell genomes that will govern all the further steps of the infection. This multistep process is governed by the targeting of viral DNA/integrase complexes, called intasomes, toward suitable chromatin regions and their subsequent functional association with nucleosomes. Several cellular factors have been shown to play a role in this targeting mechanism, such as LEDGF/p75 and CPSF6 for lentiviruses (Kvaratskhelia et al. 2014; Sowd et al. 2016) and BET proteins for gammaretroviruses (Sharma et al. 2013; De Rijck et al. 2013). Nevertheless, the integrase protein (IN) by itself has been shown to be a key determinant of integration site selection and to interact directly with chromatin components such as remodeling factors and histones (Lesbats et al. 2011; Matysiak et al. 2017; Benleulmi et al. 2017; Maskell et al. 2015). Furthermore, it has been reported that the chromatin structure surrounding the insertion site modulates retroviral integration into nucleosomes (Benleulmi et al. 2015). In particular, chromatin compaction was reported to prevent HIV-1 integration, while chromatin remodeling promoted viral DNA insertion, suggesting that additional contacts between incoming intasomes and nucleosomal substrates must be established for optimal integration(Lesbats et al. 2011; Benleulmi et al. 2015; Matysiak et al. 2017). Interestingly, the carboxy-terminal domain (CTD) of HIV-1 IN (residues 220-270) displays a strong structural similarity to Tudor domains (Lodi et al. 1995) which are known to bind histone tails (Lu and Wang 2013; Adams-Cioaba et al. 2010). These data make histone tails good candidates for potential receptors of incoming intasomes. This idea is further supported by findings showing that retroviral IN can bind, *via* its CTD directly to histone tails (Benleulmi et al. 2017; Mauro et al. 2019; Maskell et al. 2015). HIV-1 IN mutagenesis studies have shown that amino acid substitutions that affect IN binding to nucleosomes and histone tails also affect viral infectivity (Benleulmi et al. 2017). Furthermore, mutations in the CTD were reported to affect insertion site selection based on chromatin density (Benleulmi et al. 2017; Demeulemeester et al. 2014). Indeed, CTD mutations such as R231A/H, which were shown to decrease IN affinity for histone tails, also decrease viral infectivity and retarget integration sites in regions with lower nucleosome density. In contrast, mutations which increased the affinity for the nucleosome, *e.g*., D253H, were shown to improve viral infectivity (Benleulmi et al. 2017). Similar results were obtained in the case of the prototype foamy virus (PFV) intasome, for which mutations impairing the interactions of the intasome with the nucleosome also impaired integration, as observed for HIV-1 (Maskell et al. 2015).

Taken together, these data suggest that retroviral INs may have intrinsic chromatin-binding properties that enable their participation, in concert with tethering cofactors, in binding to the final chromatin insertion site. In parallel, the role of LEDGF/p75 in targeting the integration complex toward chromatin has been clearly demonstrated extensively in the literature. Furthermore, LEDGF/p75 has been shown to stimulate *in vitro* HIV-1 integration onto nucleosomes (Botbol et al. 2008), and the binding of the LEDGF/p75 PWWP domain to mononucleosomes and its preference for H3K36me3 have been reported *in vitro* (Botbol et al. 2008; Eidahl et al. 2013). However, the molecular basis for how LEDGF/p75 engages chromatin in the context of its complex with IN and how IN chromatin-binding properties may influence this targeting remain mainly unclear. The lack of experimental systems allowing monitoring of the direct association between proteins and chromatin in the absence of other cofactors prevented further characterization of these molecular processes.

Using both biochemical approaches and a chromosome-binding assay developed in this work, we investigated the LEDGF/p75-mediated chromatin tethering of the HIV-1 IN. Our study revealed that this retroviral IN has an unedited intrinsic property enabling it to bind and recognize specific chromatin regions even in the absence of its cofactor. We also found that this function is carried in the IN CTD. Furthermore, we show here that this IN chromatin-binding property is modulated by the IN interaction with LEDGF/p75, which modifies the efficiency of IN binding to chromosomes and redirects the enzyme to alternative/different chromatin regions. Finally, we depicted the chronology of the LEDGF/p75-mediated targeting of HIV-1 IN to chromatin, shedding light on the mechanism of action of LEDGF/p75 during integration.

## RESULTS

### Monitoring chromatin binding using chromosome spreads

To monitor the direct binding of protein onto chromatin, we used chromosomes isolated from blood cells and spread on glass (see **S1** and a full description in the methods section). Chromosome spreads provide important sources of biological material in which individual chromosomes can easily be identified. In metaphase chromatin structures, little to no transcription occurs^18^, demonstrating that the cellular machinery has been mainly eliminated and not available to interfere with the direct binding of our proteins of interest to complicate the study of the intrinsic chromatin-binding properties of proteins. This choice also removes a major source of variability between non-metaphasic chromosomes, which restrain the analysis to the intrinsic chromatin-binding properties of our candidates. We thus first adapted the (pro)metaphase chromosome spreading procedure described previously^19^ to maintain chromatin and nucleosome integrity and deplete the material associated with chromatin-binding proteins. For this purpose, we used synchronized peripheral T lymphocytes containing HIV-1-susceptible cells and a fixation buffer adjusted to 2/1 v/v methanol/acetic acid to maintain chromosome integrity for further binding assays. Then, the chromosomes were treated with a detergent-containing permeation buffer allowing the smooth release of the proteins associated with the chromatin except for the nucleosome components. The integrity of the chromosome was then checked by DAPI staining and immunostaining using antibodies directed against histones as previously shown^19^ (**S2**). We also ensured the absence of contamination with chromatin-associated proteins, which would have biased our analysis. In particular, we controlled for the lack of cellular factors previously reported to modulate HIV-1 integration, including FACT and LEDGF/p75, and immunostaining did not reveal these factors, confirming that they were not present, at least not in significant amounts (**S2**).

The methodology for analyzing the chromatin-binding property of a protein candidate was then established as described in **S1**. The protein of interest was incubated with the chromosome spreads under conditions described in the materials and methods section. The interaction with chromosomes was then monitored by immunofluorescence using antibodies directed against the studied factor, and the chromatin was visualized by DAPI staining. The protein interaction was evaluated after quantification of the fluorescence signal among selected chromosomes with ImageJ software. The correlation between the DAPI stained chromatin and the protein distribution was determined using a MATLAB pipeline developed in-house for this work (see materials and methods section). Briefly, DAPI staining was used to determine the exact shape and size of the chromosome (identified by inverted DAPI staining) and the distribution profile of the protein fluorescence signal along each selected chromosome. The intensity of binding, as well as the distribution of the protein, were then compiled from at least ten independent spreads and up to ten chromosomes that could be unambiguously identified and oriented by inverted DAPI staining (mainly chromosomes 1, 2 and 3). The means and standard deviations of the data were calculated. Using this method, the data on both the efficiency of the interaction and the preference for chromosomal regions were assessed by measuring the fluorescence signal along the different chromosomes.

This assay is applied herein to study the chromatin-binding property of two protein partners involved in HIV-1 integration: viral IN and cellular LEDGF/p75 proteins, which that are known to interact during integration targeting of chromosomal insertion loci.

### Analysis of intrinsic HIV-1 IN and LEDGF/p75 chromatin-binding properties

The chromatin-binding properties of LEDGF/p75 and HIV-1 IN were first analyzed independently. When incubated separately with metaphase chromosomes, both proteins showed chromatin binding. However, significant differences were detected in their preference for specific chromosomal regions. Indeed, as reported in **Figure 1A**, LEDGF/p75 showed, as expected, strong binding to chromatin.

**Figure 1.**
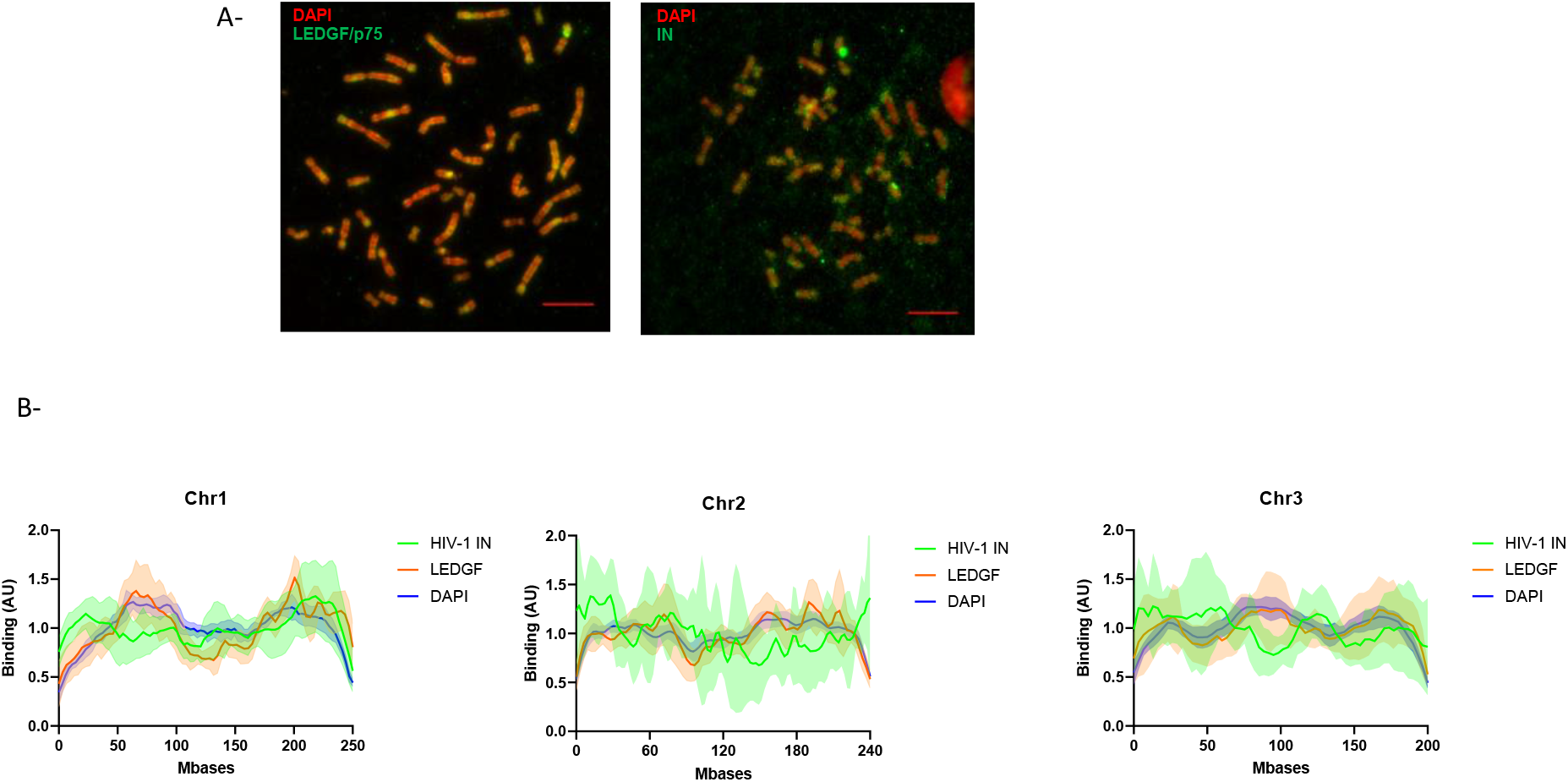
Binding of human LEDGF/p75 and HIV-1 IN to chromosomes spreads. Either recombinant purified LEDGF/p75 or HIV-1 IN (4nM) was incubated with chromosomes spreads under the conditions described in materials and methods section. Monitoring of their interaction with the chromatin was performed by immunofluorescence with the corresponding antibody coupled to ALEXA 594 (green signal) and DAPI staining (red signal) (**A**). After analysis of the signal, distribution profile of each protein was reported for each position of the chromosomes. Examples for chromosome 1, 2 and 3 are reported in (**B**). The data are reported as means from the quantification of 7 to 11 chromosomes in each condition ± standard deviation (SD). Scale bar=10μM.

An analysis of the LEDGF/p75 distribution was then performed on chromosomes unambiguously identified and oriented on the basis of DAPI staining, such as chromosomes 1, 2 and 3. The results showed that similar regions of each chromosome were apparently enriched with the factor (**Figure 1B**). LEDGF/p75 localization and DAPI staining were then compared, and statistical analysis, including Pearson correlation and p-value (percentage of comparisons with high correlations) calculation, was applied to determine the similarities and the differences between the distribution curves. The mean and standard deviation of the Pearson correlation calculation (>0 for a correlation, =0 for no correlation and <0 for a negative correlation) performed between all paired LEDGF/p75 and DAPI curves unambiguously showed a high correlation (Pearson correlation for chromosome 1 = 0.53±0.21 and p<0.01 for 84% of pairs, for chromosome 2 = 0.51+-0.18 and p<0.01 for 90% of pairs, for chromosome 3 = 0.39±0.22 and p<0.01 for 92% of pairs). Analysis of HIV-1 IN chromatin-binding properties also revealed a reproducible interaction of the retroviral enzyme with specific regions of the chromosomes (**Figure 1A**, right panel). Regions bound by IN in the context of a specific chromosome were reproducibly enriched with the retroviral enzyme (see the cases of chromosomes 1, 2 and chromosome 3 in **Figure 1B**). Strikingly, in contrast to the observation for LEDGF/p75, the comparison between IN localization and DAPI staining showed only poor, or no, correlation (Pearson correlation for chromosome 1=0.14±0.21, p<0.01 for 29% of pairs, for chromosome 2= 0.16+-0.26, p<0.01 for 50% of pairs, and for chromosome 3=0.083±0.22, p<0.01 for 15% of pairs). Merging of the LEDGF/p75 and IN chromatin interaction signal confirmed that the distribution profile of each protein showed only partial overlap (merged curved from chromosomes 1, 2 and 3 in **figure 1B**). Direct statistical comparison of the distribution of HIV-1 IN and LEDGF/p75 showed no or low correlation between these proteins on chromosome 1 (Pearson correlation=0.19±0.3, p<0.01 for 57% of pairs), chromosome 2 (Pearson correlation= 0.09+-0.24, p<0.01 for 40%) and chromosome 3 (Pearson correlation=0.04±0.31, p<0.01 for 32% of pairs). These data confirmed that both proteins bind chromatin but showed different affinities for chromosome regions.

### LEDGF/p75 modulates both the efficiency and specificity of IN binding to chromatin

In HIV-1-infected cells, IN and LEDGF/p75 are assumed to form a complex that allows efficient targeting of the intasome toward targeted chromatin sites. Thus, to investigate the impact of LEDGF/p75 on the intrinsic chromatin-binding property of HIV-1 IN, we compared the localization of IN alone or in the presence of LEDGF/p75. As reported in **Figure 2A**, IN was found to strongly bind the chromosome in the presence of its cofactor LEDGF/p75.

**Figure 2.**
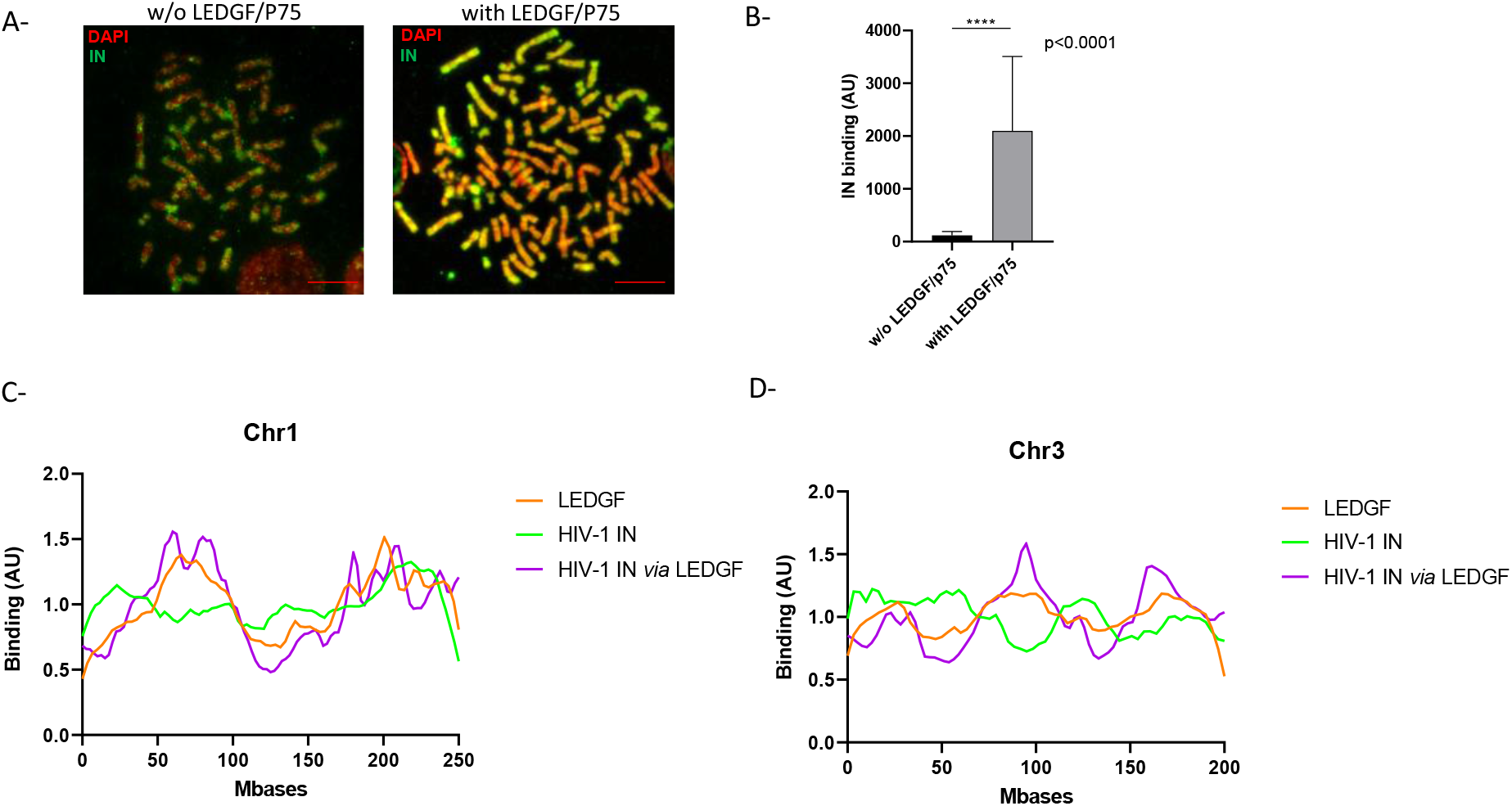
Comparison between the chromatin binding property of HIV-1 IN alone or in the presence of LEDGF/p75. HIV-1 IN was incubated with chromosomes spread in the absence or in the presence of LEDGF/p75 and its binding to chromatin was monitored by immunofluorescence with anti-IN antibodies (**A**). The intensity of total IF signal has been quantified in each conditions using ImageJ software. The data are reported as mean from the quantification of 7 to 11 chromosomes in each condition ± standard deviation (SD), p-value was calculated by student test. The distribution profile of IN, LEDGF/p75 alone or incubated together along chromosome 1 and 3 was analyzed and reported respectively in (**C**) and (**D**). The data are reported as means from 7 to 11 serials of quantifications ± standard deviation (SD). Scale bar=10μM.

Quantification of the intensity of binding of IN to chromatin in each condition (total IN fluorescence signal) along the chromosomes analyzed from the 7-11 spreads showed that the presence of LEDGF/p75 strongly and significantly improved the amount of IN bound to chromatin (see quantification data in **Figure 2B**). The analysis of the distribution of IN alone or in the presence of LEDGF/p75 was then performed and reported for chromosomes 1 and 3 in **Figure 2** panels **C** and **D**. The global distribution of IN along the chromosomes was found modified by the presence of its cofactor. Indeed, when the correlation of IN alone and LEDGF/p75 alone was compared to the correlation of IN in the presence of LEDGF/p75 and LEDGF/p75 alone the Pearson correlation between the independent distribution of IN and LEDGF/p75 shifted from 0.19±0.3, p<0.01=0.57, to 0.45±0.16, p<0.01=0.86, in the presence of LEDGF/p75 for chromosome 1 and from 0.04±0.31, p<0.01=0.32, without LEDGF/p75 to 0.23±0.25, p<0.01=0.37, in the presence of LEDGF/p75 for chromosome 3. A calculation of the p-value (Wilcoxon rank sum test) performed in each position of the chromosome (see **S3**) allowed us to more precisely define several regions of the chromosomes preferred or excluded in each condition. These statistical and correlative analyses of the IN distribution showed that in the presence of LEDGF/p75, HIV-1 IN was found to be mainly retargeted toward alternative regions of chromatin that were mainly similar to those recognized by LEDGF/p75 alone.

Taken together these data indicate that LEDGF/p75 modified IN localization along chromosomes demonstrating its IN retargeting function in our model and its capability to modulate IN chromatin binding properties.

### LEDGF/p75 modulates the efficiency but not the selectivity of the *in vitro* integration onto mononucleosomes catalyzed by IN

To further investigate the effect of LEDGF/p75 on functional integration complexes and determine whether the LEDGF/p75-mediated increase in IN chromatin-binding was related to the nucleosomal features of chromosomes, we performed typical *in vitro* integration assays using biotinylated mononucleosomes (MNs) immobilized on streptavidin beads as previously performed (Benleulmi et al. 2017; Mauro et al. 2019). As shown in **Figure 3A**, LEDGF/p75 strongly stimulated HIV-1 IN activity on MNs, while integration onto naked DNA was only slightly improved in the presence of the cofactor. These data were in agreement with data from the literature (Botbol et al. 2008) and suggested that LEDGF/p75 modulated the functional interfaces between the HIV-1 intasome and the nucleosome.

**Figure 3.**
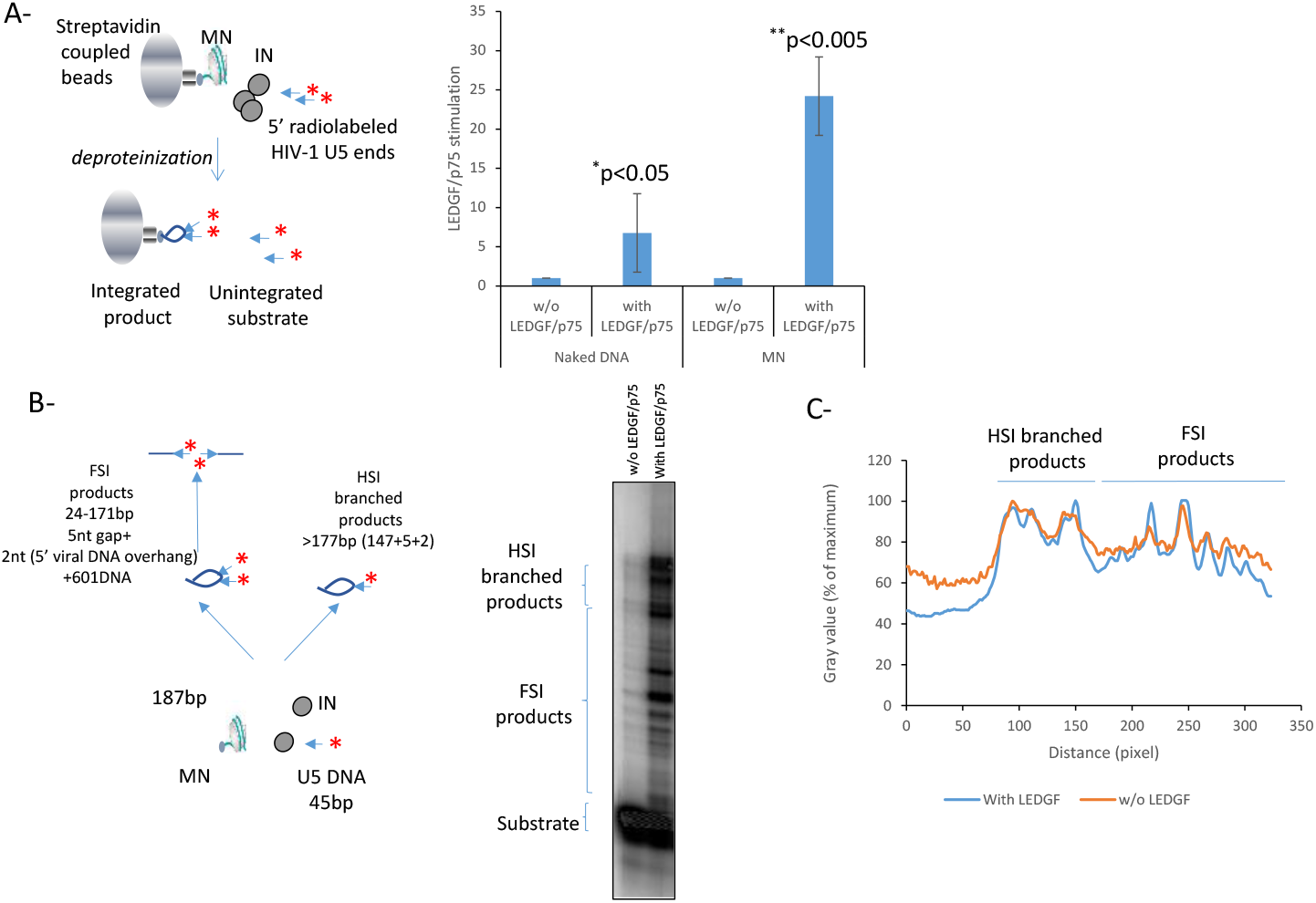
Effect of LEDGF/P75 on *in vitro* integration catalyzed by HIV-1 IN onto mononucleosomes. Integration was performed using radiolabelled viral DNA, HIV-1 IN with or without LEDGF/p75 using either biotinylated naked 601 DNA or MN assembled on 601 sequence. The integration was quantified by measuring the remaining radioactivity on streptavidin beads after hammering the target DNA (**A**) or by loading the integration products on polyacrylamid 6-12% gradient gel (**B**). Data are reported in (**A**) as stimulation effect of LEDGF/p75 according to the control experiment performed without the cellular factor on either naked of MN targets. Data were obtained from at least three independent experiments and are reported as means ±SD. Statistics were performed by student test and the calculated p is reported on the figure. Integration product structures were analyzed on native polyacrylamide 6-12% gel as reported in (**B**). The half site integration (HSI) and full site integration (FSI) products are schematized in the figure. The positions of the integration sites on the nucleosomal DNA in the absence or in the presence of LEDGF/p75 were compared by scanning the gel migration profile using ImageJ software and reporting the radioactivity signal intensity in function of the migration distance from the top of the gel (representative analysis reported in (**C**)).

We next analyzed the structure of the integration products generated on MNs using native polyacrylamide gels. As shown in **Figure 3B**, the integration products formed in the presence of IN alone were distributed along the MNs with preferred sites. While the stimulation of the integration by LEDGF/p75 was confirmed, the distribution profiles of the integration sites did not appear drastically changed, as indicated by the insertion site position analysis shown in **Figure 3C**. These results suggest that LEDGF/p75 improves nucleosomal integration catalyzed by IN without affecting the global anchoring position for the integration complex onto MN. Taken together, these biochemical data agree closely with the findings showing the stimulation of IN binding to chromatin by LEDGF/p75 observed on chromosomes (cf **Figure 3A**), indicating that the cellular factor potentiates IN functional interactions with nucleosomes and modulate the chromatin binding properties of the retroviral enzyme. The similar integration profile observed with *in vitro* integration assays additionally suggests that the differences observed in IN binding to chromatin in the presence of LEDGF/p75 were not due to altered specificity for the nucleosome structure but were mainly related to distinct preference for local surrounding chromatin features, suggesting a role of chromatin neighboring the interaction site.

### HIV-1 IN and LEDGF/p75 determinants of chromatin tethering and targeting processes

The LEDGF/p75-IN interaction sites have been precisely mapped on both partners, and the IN-binding domain (IBD) of LEDGF/p75 and the catalytic core domain of IN (CCD) have been identified as interacting regions (see **Figure 4A**). The IBD D366N mutation has been shown to prevent the ability of LEDGF/p75 to interact with IN (Cherepanov 2007), and extensive studies have shown that the PWWP domain and AT hook domains of LEDGF/p75 mediate its chromatin-binding properties (Eidahl et al. 2013; Llano et al. 2006; Shun et al. 2008). Analysis of the D366N mutant and hook-deleted PWWP/AT constructs allowed us to dissect the LEDGF/p75-mediated anchoring of IN to chromatin. Specifically, we showed that the D366N mutation is associated with a decrease in the capability of LEDGF/p75 to target IN toward chromatin (**Figure 4B**, quantification data are in **4C**). Furthermore, deletion of the PWWP and AT hook domains (mutant 208-440 and mutant Δ141, respectively) prevented IN chromatin targeting by LEDGF/p75 (**Figure 4B-C**).

**Figure 4.**
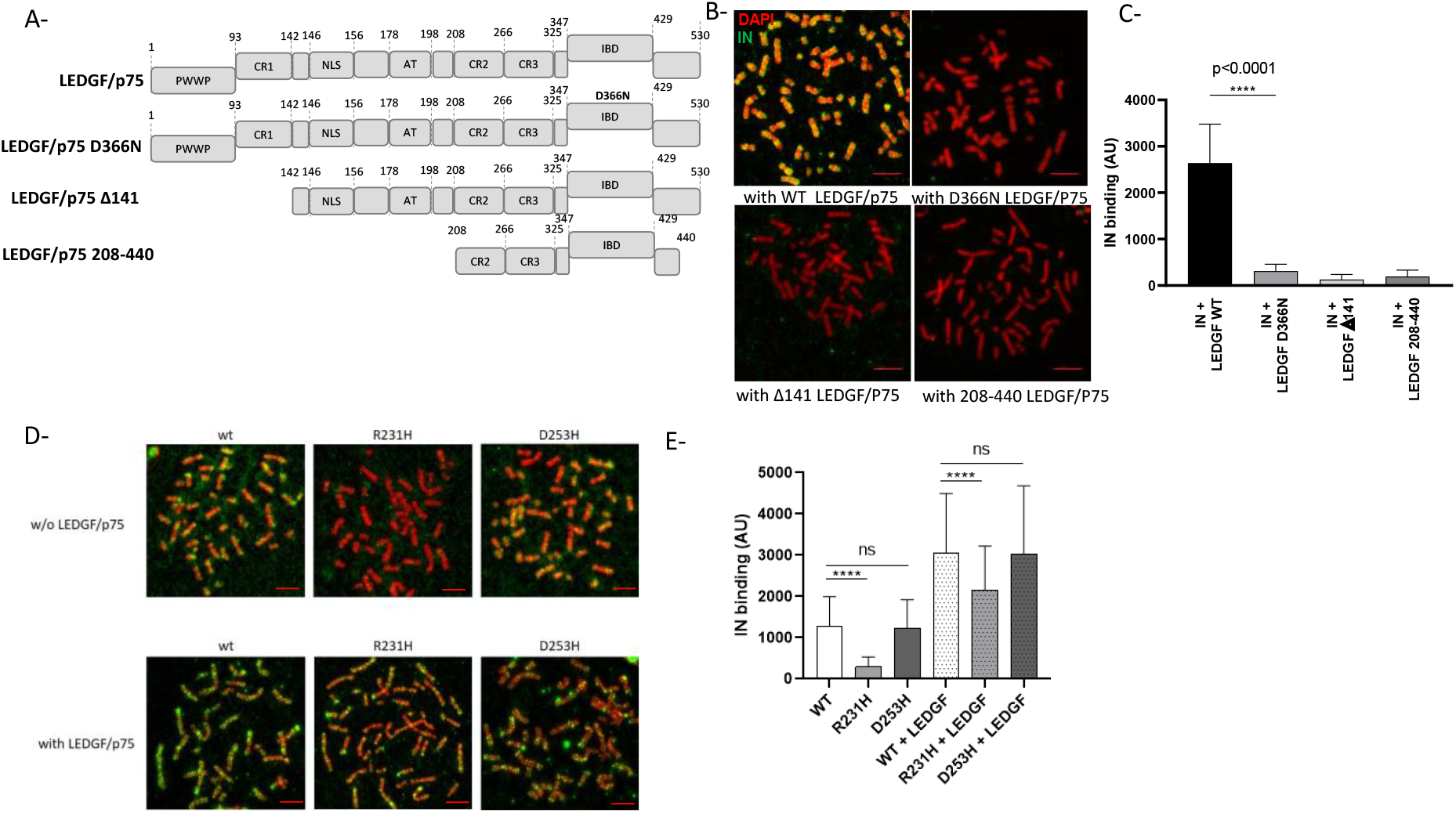
Effect of LEDGF/p75 and IN carboxyterminal domain mutants on the modulation of HIV-1 IN chromatin binding property. The different constructions used in the experiments are reported in (**A**). Full length, D366N LEDGF/p75 punctual mutant impaired for binding to HIV-1 IN and truncations deleted for the PWWP chromatin binding domain and AT hook of LEDGF/p75 were compared for their effect on HIV-1 IN binding to chromosomes. The purified LEDGF/p75 mutants were added to HIV-1 IN in the chromatin binding assay and theirs effects on the anchoring of IN were compared to the one showed by WT protein (**B**). The total IN binding signal was quantified along chromosomes (here chromosome 1) in the different conditions and are reported in the histogram as means of eight independent set of quantification (between 738 to 954 quantification points) ± standard deviation (SD). Statistics were performed by student test and the calculated p is reported on the figure (**C**). WT, R231H and D253D IN proteins were incubated with chromosome spreads in the absence or in the presence of LEDGF/p75 (4nM) and the IN binding to chromatin was monitored by immunofluorescence using anti-IN antibody (**D**). The intensity of total IF signal has been quantified in each condition using ImageJ software on several chromosomes and reported as IN binding. The data are reported as mean of the quantification of ten chromosomes 1 (1026-1122 quantification points) ± standard deviation (SD). Statistics were performed by student test and the calculated p is reported on the figure (**E**). Scale bar=10μM.

Since IN alone shows affinity for chromatin, we also investigated the molecular determinants of this interaction and their roles in tethering the viral enzyme in the presence of LEDGF/p75 to chromatin. Target DNA and histone-binding properties have been described for the CTD of several retroviral INs, including HIV-1 IN (Maskell et al. 2015; Benleulmi et al. 2017; Mauro et al. 2019). Thus, we analyzed the importance of this domain in binding to chromosomes. Several amino acids have been identified in HIV-1 IN as important for the interaction to both the target DNA and histone tail (Benleulmi et al. 2017; Mauro et al. 2019; Serrao et al. 2014; Demeulemeester et al. 2014). These positions include the R231 and D253 residues. To test the chromatin-binding properties of HIV-1 IN, we used recombinant INs mutated in these positions and showed that these mutations affected their association with nucleosomes and histone tails (see extended analysis of the CTD mutants in (Benleulmi et al. 2017) and in Figure **S4**). As reported in **Figure 4D** and quantified in **4E**, the R231H mutant IN was found to poorly bind chromatin, correlating closely with its poor ability to bind nucleosomes *in vitro*. In contrast, D253H mutant of IN, which presented similar, or even better, affinity for nucleosomal DNA than the WT enzyme, bound chromatin with efficiency close to WT. We next investigated whether this intrinsic chromatin-binding property of the IN CTD also contributes to the interaction of the viral protein in the presence of LEDGF/p75 with the chromosome. The R231H and D253H IN proteins binding to chromosome were next analyzed in the presence of LEDGF/p75. In this context, the differences found in the association of the mutants with chromatin compared to that of the wild-type enzyme were drastically smoothed, confirming that LEDGF/P75 is a strong modulator of the chromatin-binding property of IN (**Figure 4D-E**). However, in the presence of LEDGF/p75, the R231H chromatin-binding efficiency was still significantly lower than that of the wild-type and D253H mutant proteins. Taken together, our results indicate that the CTD domain of HIV-1 IN participates in the recognition and binding of the viral protein to the chromatin regions of the chromosomes, even in the presence of its cofactor.

### Optimal IN targeting toward chromatin requires preformed IN-LEDGF/p75 complex

Since the chromatin binding assay (CBAssay) performed in this work recapitulated the LEDGF/p75 targeting of IN toward chromatin, we used this model to depict the targeting process better. The time-of-addition experimental scheme presented in **Figure 5A** shows that, although incubation of LEDGF/p75 with chromosomes before adding IN (i) slightly improved IN binding to chromatin, the best IN chromatin anchoring by LEDGF/p75 was observed when both proteins were preincubated before their addition to the chromosome spreads (**Figure 5B**, condition (ii)).

**Figure 5.**
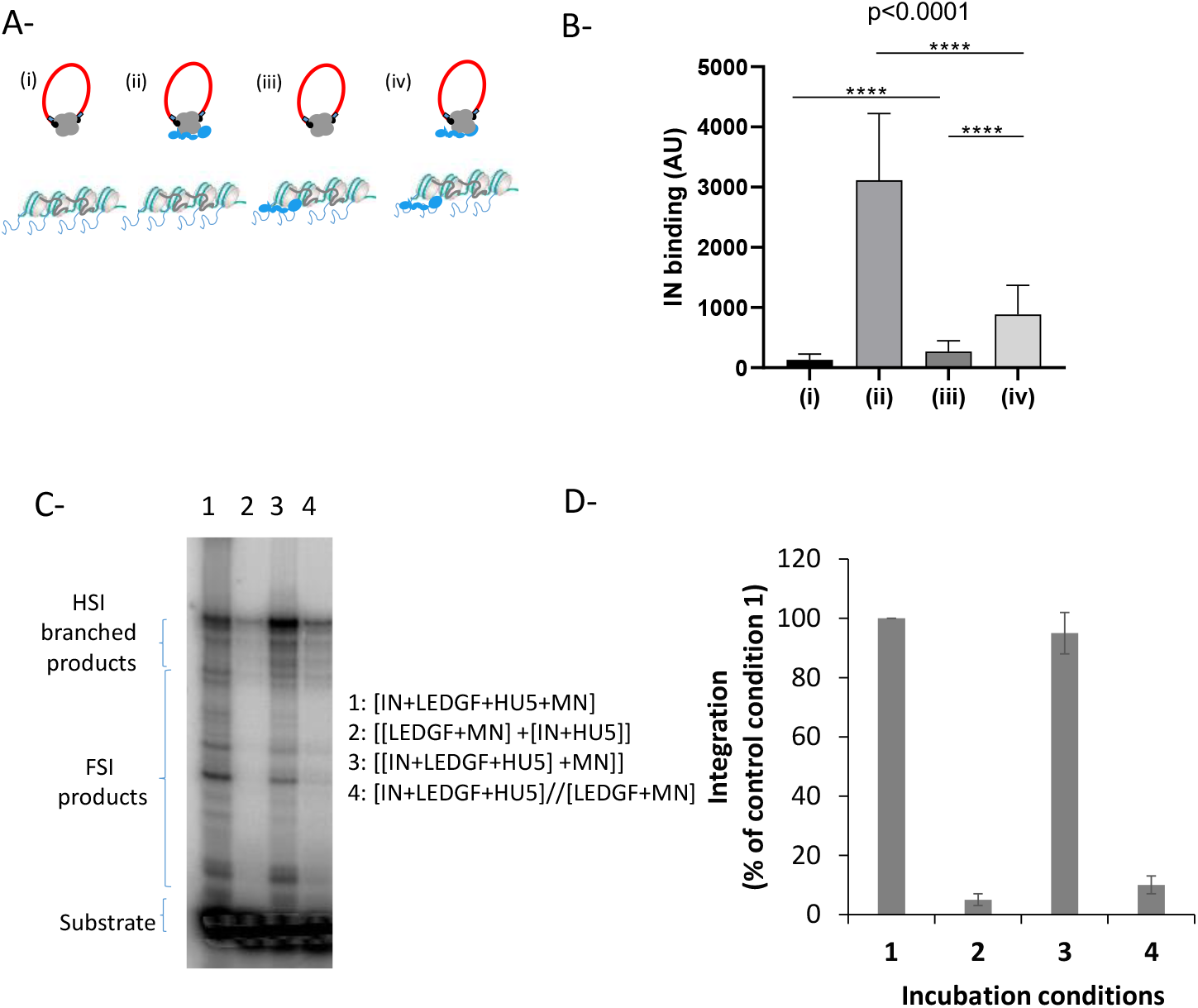
Time of addition analysis of the LEDGF/p75-mediated chromatin tethering of HIV-1 IN. HIV-1 IN was incubated with chromosome spreads in the presence or in the absence of LEDGF/p75 following different time of addition conditions reported in (**A**): (i) IN alone, (ii) IN and LEDGF/p75 were preincubated before being added to the chromosome spreads, (iii) LEDGF/p75 was pre-incubated with chromosome spreads before adding IN and (iv) IN and LEDGF/p75 were preincubated together and added to chromosomes spreads previously precincubated with LEDGF/p75. The total amount of LEDGF was similar in all conditions. The intensity of total IN IF signal has been quantified in each condition using ImageJ software on several chromosomes and reported as IN binding. The data are reported in (**B**) as mean of the quantification from 12 chromosomes (here chromosome 3, 1135-1220 quantification points per condition) ± standard deviation (SD). Statistics were performed by student test and the calculated p is reported on the figure. Integration assay onto MN was performed in difference incubation conditions using recombinant IN with or without LEDGF/p75 as done in **Figure 4B**. A representative gel is reported in (**C**) and quantification data obtained from at least three independent experiments are reported IN in the graphic (**D**) as means ±SD. Integration product structures are reported as half site integration (HSI) or full site integration (FSI). Condition 1: IN, LEDGF/p75, viral donor U5 DNA and target nucleosomal DNA were pre-incubated altogether, condition 2: [LEDGF/p75/target nucleosomal DNA] and [IN/viral U5 donor DNA] were preincubated separately before reaction, condition 3: IN, LEDGF/p75 and the viral donor U5 DNA were preincubated together before adding the target nucleosomal DNA, condition 4: [IN, LEDGF/p75, viral donor U5 DNA] and [LEDGF/p75, target nucleosomal DNA] were incubated separately before reaction.

Interestingly, preincubation of LEDGF/p75 with chromosomes before adding the preformed IN•LEDGF/p75 complex (conditions (iii) and (iv)) decreased the IN-binding efficiency compared to conditions without LEDGF/p75 bound to chromosomes. These data suggest, first, that the efficient retargeting of the incoming IN requires the formation of the complex before interacting with chromatin, and second, the presence of LEDGF/p75 on the final anchoring sites may prevent optimal binding. We further investigated this process with a functional integration assay to determine whether the integration process followed similar chronological rules. As reported in **Figure 5C** and quantification in **5D**, preincubation of IN with LEDGF/p75 led to maximal integration efficiency *in vitro*, while the incubation of LEDGF/p75 with nucleosomes before adding IN or the preformed IN•LEDGF/p75 complex profoundly prevented optimal integration. These data are in agreement with the chromatin binding of IN observed in the CBAssay, supporting the functional relevance of our system and strengthening our conclusions.

Altogether, these results indicate that without preliminary interaction between both factors, IN binding to chromatin was relatively inefficient, and IN alone does not appear efficient in finding LEDGF/p75-enriched sites on chromosomes. In addition to the data obtained in the integration assays, this finding further supports the hypothesis that incoming IN forms a functional complex with LEDGF/p75 before reaching chromatin sites. We further investigated this finding by analyzing the behavior of fully functionally pre-assembled retroviral intasomes.

### Chromatin-binding properties of functional intasomes

The CB assay was applied to analyze and compare the chromatin-binding properties of functional intasomes. We used the maedi-visna virus (MVV) intasome as a lentiviral model constructed following reported procedures, allowing us to obtain sufficient amounts of complex suitable for binding analysis, in contrast to the HIV-1 intasome (Passos et al. 2017; Ballandras-Colas et al. 2017). The MVV intasome was assembled with recombinant MVV IN and LEDGF/p75 proteins and short ODN containing a viral end DNA sequence tagged with FITC. This intasome checked in typical *in vitro* integration assays and was found to be catalytically competent (see the purification and integration assays in **S5**). This purified functional complex was thus analyzed with our CBAssay. As shown in **Figure 6A**, the MVV intasome was found to efficiently bind chromosomes with preference for some chromatin regions (see an example of the distribution profile on chromosome 1 in **Figure 6C**).

**Figure 6.**
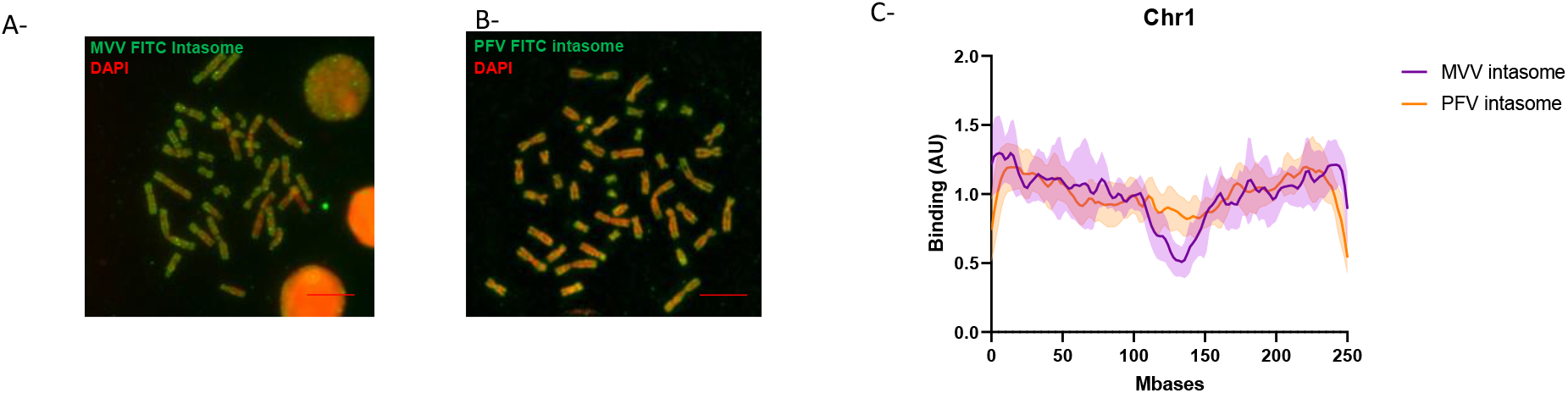
Analysis of the chromatin binding property of MVV and PFV intasomes. Each intasome was assembled using the corresponding recombinant protein (IN and LEDGF/p75 in the case of the lentiviral MVV complex), the viral short DNA sequence corresponding to the viral end tagged with FITC. The intasomes were purified by size exclusion chromatohraphy (see **S5**). After verification of their functionality in *in vitro* concerted integration assays (**S5**) the intasome (4nM) were incubated with chromosomes spreads. Interaction profile was determine by FITC-epifluorescence acquisitions (MVV intasome in **A**, and PFV intasome in **B**). The distribution profile of each intasome onto chromosome 1 was compared in (**C**). The data are reported as means from the quantification of eight to nine chromosomes ± standard deviation (SD). Scale bar=10μM.

To determine whether this profile was specific for each retroviral intasome, we analyzed the chromatin-binding properties of an additional functional intasome: the prototype foamy virus (PFV). Following indications from the literature (Maskell et al. 2015), we assembled a PFV intasome exhibiting the proper integration activity (**S5**). Analysis of the chromatin-binding properties of the purified intasome as determined with the CB assay also indicated that this complex binds chromatin (**Figure 6B**). However, as shown in **Figure 6C**, the comparison between the MVV and PFV intasome distribution profiles showed only a weak correlation (Pearson correlation= 0.28±0.20, p<0.01=0.55), and a statistical correlation analysis performed with chromosome 1 showed a significant absence of exclusion from the centromeric region, as had been observed for the MVV intasome (**Figure 6C** and **S6**). Indeed, the PFV intasome chromatin binding was more homogeneous than for MVV intasome or HIV-1 IN•LEDGF/p75 complex. A control experiment performed with PFV in the presence of LEDGF/p75 showed no effect of the cellular factor, as expected from the specificity of the cellular protein for lentiviral INs (Figure **S6**). This result confirmed that the differences found between the two intasomes were not due to LEDGF/p75 but to intrinsic chromatin-binding properties specific for each complex, further validating the specificity of our assay. Taken together, our data demonstrate that the structure of the integration complex and the presence of cellular tethering factors are major determinants of chromatin preference.

## DISCUSSION

During the retroviral integration process, the association between the incoming intasome and host chromatin constitutes a point of no return and a platform for molecular exchanges between the virus and the cell. While the early targeting processes of the intasome toward chromatin have been investigated for a long time, its final association with the nucleosome is not well characterized. Recent advances have shown that after nuclear entry and targeting toward a suitable region of the genome by cellular IN or CA cofactors, such as CPSF6 and LEDGF for lentiviruses, the intasome and the histone component of the nucleosome directly interact (Maskell et al. 2015; Benleulmi et al. 2017; Mauro et al. 2019). Furthermore, IN domains, such as the CTD, have been shown to directly engage the target DNA and participate in insertion site selection in chromatin (Serrao et al. 2014; Demeulemeester et al. 2014). These data indicate that the CTD of INs is the central domain for the association of IN with the target substrate via its direct DNA- and histone-binding properties. In the specific case of HIV-1, previous data showed that HIV-1 IN association with histone tails, especially the H4 tail, modulated its functional binding to the nucleosome, leading to structural changes and the stabilization of the intasome/nucleosome complex (Benleulmi et al. 2017; Mauro et al. 2019). Mutations within the CTD that affect retroviral IN-histone binding were also shown to affect both the functional association with the nucleosome integration site selectivity *in vitro* and *in vivo* (Maskell et al. 2015; Benleulmi et al. 2017). Taken together, these data support the presence of a chromatin-binding function in the CTD of IN that may participate in the functional association with chromatin and insertion site selection. However, to date, no model for validating this function or investigating its molecular determinants is available. Here, we established a system allowing us to directly monitor the chromatin-binding property of proteins, and we applied this model to the study of factors important for HIV-1 integration. Using this system, we were able to show the chromatin-binding properties of functional retroviral intasomes and their cellular and viral components.

The analysis of the chromatin-binding property of HIV-1 IN alone showed that the protein was able to selectively bind regions of chromatin with certain specificity. Since no endogenous FACT or LEDGF/p75 was detected (cf **S2**), here, IN binding to chromatin was more likely an intrinsic property of the retroviral enzyme and not due to contamination by IN cofactors, which have the potential to interact both with chromatin and IN. Both the specificity and efficiency of IN binding to chromosomes were affected by the presence of the IN cofactor LEDGF/p75. A similar integration profile observed in integration assays performed on mononucleosomes *in vitro* additionally suggested that the differences observed in IN binding to chromatin in the presence of LEDGF/p75 were not due to altered nucleosome specificity but were mainly related to distinct preferences of the IN•LEDGF/p75 complex for local chromatin features. This outcome suggests a role for nucleosomes neighboring the final interaction site selected within chromatin. The analysis of the LEDGF/p75 distribution along the chromosomes showed that it appeared to correlate closely with chromatin DAPI staining. In contrast, the poor correlation found between IN distribution and DAPI staining suggested that the chromatin features recognized by IN were mainly independent of DNA structure and composition. While the exact nature of the recognized chromatin regions remains to be fully elucidated, our data confirmed that the intrinsic HIV-1 IN protein chromatin-binding properties were modulated by its cofactor, LEDGF/p75. The intrinsic chromatin-binding function of IN appeared to be carried by its CTD since mutations in this domain affected the efficiency of its chromosome interaction. In particular, mutations that were shown to affect the interaction of the protein with nucleosomal DNA and histone tails (Benleulmi et al. 2017; Mauro et al. 2019) affected the interaction with chromosomes in a similar manner. These results correlate closely with cellular data previously reported, *i.e*., a decrease in the integration efficiency of the R231H mutant and a close or better integration efficiency of the D253H mutant, with a redistribution of integration sites in both cases (Benleulmi et al. 2017). Indeed, the efficiency of cellular integration is directly related to the efficient association between the incoming integration complex, as shown in the CBAssay, while the selection of the final integration site may depend in part on the intrinsic capability of the enzyme to prefer chromatin structures even in the presence of LEDGF/p75, as demonstrated here.

Furthermore, comparisons of both the IN distribution profile and chromatin-binding efficiency in the presence or absence of LEDGF/P75 clearly showed that cellular factors modulate IN binding properties. Indeed, LEDGF/p75 greatly enhanced IN binding to chromosomes and modified its localization, targeting it mostly toward LEDGF/P75-preferred sites. We showed that both the PWWP chromatin binding and IN-binding domains in LEDGF/p75 were required for IN targeting toward chromatin. Our data indicated that LEDGF potentiated the IN functional interaction with nucleosomes and, thus, modulated the retroviral enzyme chromatin-binding property by affecting its behavior on nucleosomes. Interestingly, the R231H mutation in the IN CTD was shown to affect the enzyme chromatin-binding property, even in the presence of LEDGF/p75, suggesting that the IN CTD may participate in the final association and insertion site selection. This may provide the molecular basis for previously reported cellular selectivity analyses of viruses carrying these CTD mutations (Benleulmi et al. 2017). Interestingly, all these data appear to corroborate of previous works showing that HIV-1 IN may modulate the interaction of LEDGF/p75 with chromatin (Astiazaran et al. 2011). Indeed, the efficiency of viral integration into cells is directly related to the efficiently established association between the incoming integration complex, as shown in the CBAssay, while the selection of the final integration site may be mediated in part by the intrinsic ability of the enzyme to prefer chromatin structures, even in the presence of LEDGF/p75, as demonstrated herein. Further characterization of the IN targeting process via LEDGF/p75 showed that maximal efficiency was observed when IN was precomplexed with its cofactor. This finding supports a model where the presence of LEDGF/p75 complexed to the incoming integration complexes allows efficient anchoring to the final insertion sites, revealing a longstanding issue in the field. This supposition was confirmed by analyzing purified functional MVV lentiviral intasomes assembled with LEDGF/p75, which showed very efficient binding to chromosomes. The comparison with the PFV intasome profile also suggested that the distribution profiles were intasome-specific, indicating that the integration complexes carry the main determinants for local chromatin anchoring through their nucleosome binding sites.

In addition to providing important mechanistic information about the integrated chromatin targeting process, the CB assay used here paves the way for broad and deep analyses of these chromatin-binding mechanisms. Indeed, data obtained with functional intasomes indicated that the chromosome spread model can recapitulate the final step of chromatin targeting by the integration complexes. Consequently, this model constitutes a suitable system for further depicting all the parameters of this mechanism, such as the roles of additional cofactors or the kinetics of the process. Additionally, the CBAssay can also be used in structure/function studies to analyze the role of specific amino acids and domains of the different integration actors in the anchoring mechanism, as shown here for the analysis of some IN CTD mutants. Testing small molecules that may affect the chromatin-binding function of the enzymes should also be possible with this assay, which may become complementary to the current panoply of biochemical and cellular assays used for studying the regulation of retroviral integration by chromatin components. Finally, our model may also be used to study any protein or nucleocomplex candidate that may have chromatin-binding properties and thus may be applied to broader fields, such as cancer or gene development.

## METHODS

### Proteins purification and antibodies

The HIV1-IN was expressed in E.coli (Rosetta) and the cells were lysed in buffer containing 50mM Hepes pH 7.5, 5mM EDTA, 1mM DTT, 1mM PMSF. The lysate was centrifuged and IN extracted from the pellet in buffer containing 1M NaCl, 50mM Hepes pH 7.5, 1mM EDTA, 1mM DTT, 7mM CHAPS. The protein was then purified on butyl column equilibrated with 50mM Hepes pH7.5, 200mM NaCl, 1M ammonium sulfate, 100mM EDTA, 1mM DTT, 7mMCHAPS, 10% glycerol. The protein was further purified on heparin column equilibrated with 50mM Hepes pH7.5, 200mM NaCl, 100mM EDTA, 1mM DTT, 7mM CHAPS, 10% glycerol. LEDGF/75 was expressed in PC2 bacteria and the cells were lysed in lysis buffer containing 20mM Tris pH 7.5, 1 M NaCl, 1mM PMSF added lysozyme and protease inhibitors. The protein was purified by nickel-affinity chromatography and the His-tag was removed with 3C protease, 4°C over night. After dilution down to 150mM NaCl, the protein was further purified on SP column equilibrated with 25mM Tris pH7.5, 150 mM NaCl (gradient from150 mM to 1M NaCl), then DTT was added to 2mM final and the protein was concentrated for Gel filtration. Gel filtration was performed on a superdex 200 column (GE Healthcare) equilibrated with 25mM Tris pH7.5, 500 mM NaCl. Two mM DTT were added to eluted protein that was then concentrated to about 10mg/ml.

Polyclonal anti-HIV-1 IN antibody was purchased from Bioproducts MD (Middletown, MD, USA). Monoclonal anti-HIS was purchased from Abcam. HRP-conjugated secondary anti-rabbit antibody was purchased from Sigma and anti-mouse from Jackson ImmunoResearch. Monoclonal Anti H4 was purchased from Abcam (ref 61521 (1/400)). Monoclonal Anti LEDGF/p75 was purchased from BDbiosciences (ref 61175 (1/600)). Polyclonal Anti-SSRP1 was purchased from Abcam (ref 21584 (1/140)) and monoclonal Anti-SSRP1 was purchased from Abcam (ref 26212 (1/400)). Polyclonal anti-Rabbit IgG Secondary Antibody, Alexa Fluor 488 was purchased from FISHER SCIENTIFIC (1/400). Polyclonal anti-Rabbit IgG Secondary Antibody, Alexa Fluor 594 was purchased from FISHER SCIENTIFIC (1/400). Polyclonal anti-Mouse IgG Secondary Antibody, Alexa Fluor 546 was purchased from FISHER SCIENTIFIC (1/400). The 5’-biotinylated 601 sequence was purchased from TEBU-Bio. Biotinylated histone tail peptides were purchased from Eurogentech (Angers, France).

### Intasomes assembly

MVV intasomes were purified essentially using the method described in(Ballandras-Colas et al. 2017) with some modifications. Briefly recombinant MVV IN (16μM) was incubated with LEDGF/p75 (8μM) and it’s cognate pre-processed vDNA (5μM) coupled to FITC at 37°C for 13mins in a buffer containing 25mM BTP pH6.5, 40mM KAc, 3mM CaCl2, 10μM ZnCl2, 2mM DTT and 80mM NaCl in a final volume of 200μL. After the incubation, the mixture was adjusted at 310mM NaCl and placed on ice for 5 minutes. Intasomes were purified with a SuperDex200 10/300 24mL equilibrated with 310mM NaCl, 25mM BTP pH6.5 and 3mM CaCl2. Intasomes fractions (eluted around 7.5mL) were pooled and concentrated with a 30kDa cut-off vivaspin column. PFV intasomes were purified essentially using the method described in(Maskell et al. 2015). Briefly, recombinant PFV IN (135μM) and it’s cognate pre-processed vDNA (80μM) labelled with FITC were dialyzed over night at 18°C against a buffer containing 20mM BTP pH7.5, 200mM NaCl, 25μM ZnCl2, 2mM DTT using a 3.5kDa cut-off dialysis cassette. The mixture was then loaded on a SuperDex200 10/300 24mL equilibrated with 320mM NaCl and 20mM BTP pH7.5. Intasomes fractions (eluted around 10.5mL) were pooled and concentrated with a 30kDa cut-off vivaspin column. The sequence of the ODN used for the intasome assembly are reported in **S7**.

### Chromosomes spreads

Chromosome spreads were obtained after culture of peripheral blood T-lymphocytes, in accordance with standard procedures used for conventional karyotyping on peripheral blood cells. Briefly, 0.5 ml of peripheral blood was added to 5 ml of specific culture medium (ChromoSynchroPTM kit, EuroClone, Pero, Italy), T-lymphocytes were thereby cultured for 72 hours. After 48 hours of culture, reagents for lymphocytes culture synchronization (ChromoSynchroPTM kit, EuroClone, Pero, Italy) were added, according to the manufacturer’s recommendations. Mitotic cells were arrested at the prometaphase stage by adding to the culture 50 μl of Colcemid at 10μg/ml (KaryoMAX ColcemidTM, Thermo Fisher Scientific, Waltham, MA, USA) for 1 hour, and next, cells were fixed by a solution of methanol and acetic acid (2/1, v/v). Finally, chromosome spreading was performed on Superfrost PlusTM slides (Thermo Fisher Scientific, Waltham, MA, USA).

### Chromosomes interaction assays

Chromosomes spreads slides wer permeabilized with Triton X-100 0.1%, saponine 0.4% for 15 min and blocked with 2% bovine serum albumin (BSA), saponine 0.1%, SVF 2% for 1 hour. All incubations were carried out at room temperature and were followed by three PBS-saponin 0.1% washes. Chromosomes spreads were incubated with IN or LEDGF diluted in blocking buffer for 1 hour. Then, chromosomes spreads were incubated with primary antibodies for 1 h and secondary antibodies for 45 min. Antibodies were diluted in blocking buffer too. DNA were stained with Dapi (100ng/ml). PBS washes then water wash are performed. Finally, chromosomes spreads were mounted onto glass slides with Prolong Diamond (Life Technologies). Epifluorescence microscopy was carried out on a Zeiss AxioImager Z1 using a 63× objective. Images were acquired in FIJI.

### Analysis of chromatin binding

The interaction with chromosome was monitored by immunofluorescence using antibodies directed against the studied factor and chromatin was monitored by the DAPI staining. The fluorescence signal among selected chromosomes was quantified with ImageJ software after substracting the background using 50 pixels rolling ball radius. The different channels were merged and a segmented line of 4 units width was drawn from both sides of the analyzed chromosomes previously identified by inverted DAPI staining and oriented by centromere positioning. The DAPI staining was used to precisely determine the exact shape and size of the chromosome and the distribution profile of the protein fluorescence signal along each chromosome. The intensity of binding, as well as the distribution of the protein, were then compiled from at least ten independent spreads and up to ten chromosomes which could be unambiguously identified. Correlation between the DAPI stained-chromatin and the protein distribution was performed using MATLAB. Means and standard deviation of the data were calculated. Pearson correlation for each pair of curves was computed and the proportion of significant correlation (p value) was also provided for each analysis. In that analysis the Pearson correlation will be >0 if correlation, =0 if no correlation, <0 if anti-correlation. The p-value indicates if the correlation (or anti-correlation) is significant. The mean correlation +/- standard deviation is provided among all the comparison and a positive mean correlation indicates that most of the time, the curve correlates together, a close to 0 mean can indicate that curves do not correlate together. The p-value was also calculated as ranksum tests at each position along the chromosome in order to identify regions where there is a significant difference while taking into account the distribution profile variability among all the measurments. In this analysis a p-value <0.01 in one region indicates there is a significant difference and a p-value >0.05 indicates no significant difference meaning ether that the compared profiles are highly the same in the 2 groups or the compared profiles are highly variable in the 2 groups.

### Integration assays

HIV-1 concerted integration assays were performed as previously reported (Benleulmi et al. 2017) using recombinant purified IN or IN•LEDGF/p75 complex (200 nM in IN monomers) and biotinylated 601 mononucleosome. Under typical conditions IN/viral DNA complexes were pre-assembled 20mn at 4°C in the presence of the 17/19 bp U5 viral ends DNA hybrid radiolabeled in 5’ (10 nM). The pre-assembled complex was then incubated with MN (50ng DNA) for 5mn in 20mM HEPES pH7, 10mM MgCl2, 20μM ZnCl2, 100mM NaCl, 10mM DTT final concentrations. After reaction products are deproteolysed proteinase K treatment and phenol/chloroform/isoamyl alcohol (25/24/1 v/v/v) treatment before salt precipitation. Reaction samples are then loaded on 6-15% native polyacrylamide gel and run for 5 hours at 200V. The gel was then dried and submitted to autoradiography before analysis using ImageJ software. For quantitative assay the acceptor MN substrates were immobilized on streptavidin-coupled beads before reaction and the reaction products were deproteinized as described above and the integration was quantified by counting the remaining radioactivity bound to magnetized beads. Integration assays with intasome were performed as followed: 200ng of mononucleosome were incubated with either PFV (30nm final concentration) or MVV (70nm final concentration) purified intasome in 100mM NaCl, 20mM BTP pH 7, 12.5mM MgSO4 in a final volume of 40μL. The mix was then incubated at 37°C for 15mins (for PFV) or 60mins (for MVV). The reaction was stopped by the addition of 5.5μL of a mix containing 5% SDS and 0.25M EDTA and deproteinized with proteinase K (Promega) for 1hour at 37°C. DNA was ethanol precipitated and integration products analyzed on a 8% non denaturing polyacrylamide gel.

## AKNOWLEDGEMENT

This work has been supported by the French ANRS research agency ECTZ115893 and SIDACTION 16-1AEQ-10465. We thank N. Landrein and M. Bonhiver (MFP laboratory) for support in microscopy imaging.

