## Supplementary material for "Analysis of the intrinsic chromatin binding property of HIV-1 integrase and its regulation by LEDGF/p75 using human chromosomes spreads": Sup data 1 to 7

**Supplementary data**

**Table of contents:**

**S1. Schematic representation and pipeline of the chromosome binding assay.**

**S2. Protein composition analysis of the chromosomes spreads.**

**S3. Statistical analysis of the distribution profile of IN alone or in the presence of LEDGF/p75 on chromosomes 1 and 3.**

**S4. Functional association of WT, D253H and R231H IN CTD mutants to mononucleosomes.**

**S5. Assembly, purification and biochemical characterization of MVV and PFV intasomes.**

**S6. Analysis of the MVV and PFV intasome binding to chromosome 1.**

**S7. Sequence of the U5 ODN used for intasome assembly.**

**S1. Schematic representation and pipeline of the chromosome binding assay.** The recombinant proteins, or assembled complexes, are incubated with chromosomes spreads obtained after culture of peripheral blood T-lymphocytes (**A**). Binding to chromosome is monitored by immunodetection using adapted antibodies and chromosomes shape is measured by DAPI staining (**B**). Both DAPI and immunofluorescence staining are quantified using ImageJ software. Raw data are extracted and distribution profiles on each chromosome are determined by MATLAB-driven matching of both signals. See the materials and methods section for the full description of the incubation, acquisition and analysis procedures.

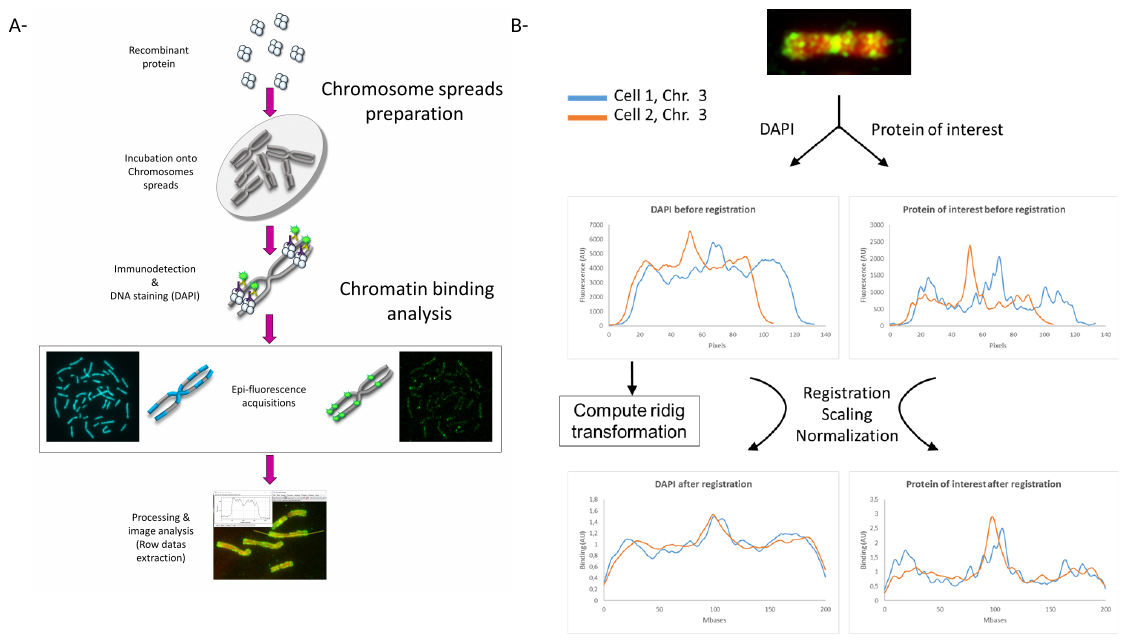

**S2. Analysis of integrity and protein composition of the chromosomes spreads.** The presence of intrinsic chromatin component histone H4, HIV-1 IN cofactors LEDGF/p75 and FACT was monitored by immunofluorescence using the corresponding antibody.

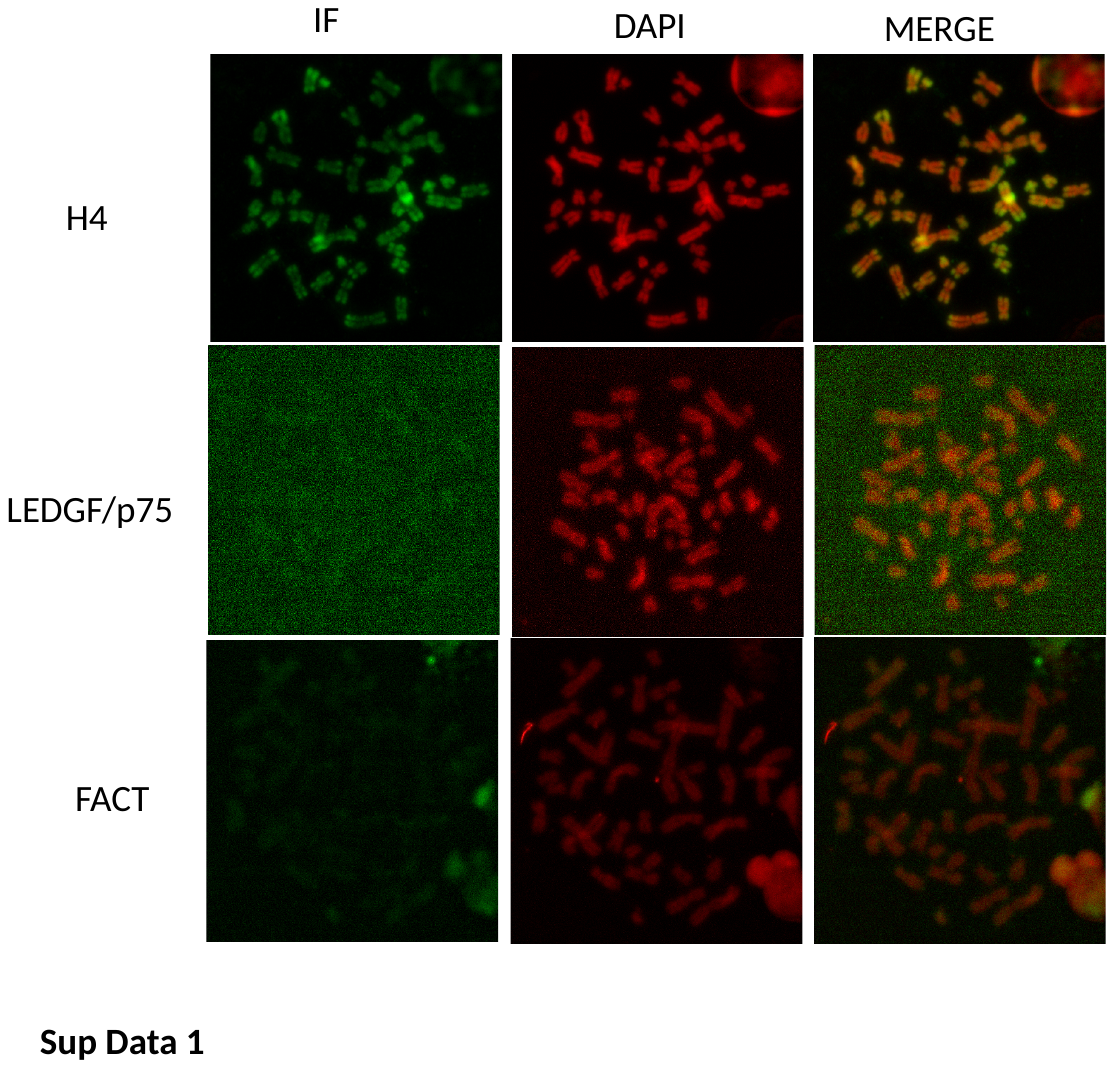

**S3. Statistical analysis of the distribution profile of IN alone or in the presence of LEDGF/p75 on chromosomes 1 and 3.** Pearson correlation for each pair of curves was computed and the proportion of significant correlation (p value) was provided for each analysis >0 if correlation, =0 if no correlation (top graph). The mean correlation +/- standard deviation is provided among all the comparison and a positive mean correlation indicates that most of the time, the curve correlates together, a close to 0 mean can indicate that curves do not correlate together. The p-value was also calculated as ranksum tests at each position along the chromosome in order to identify regions where there is a significant difference while taking into account the distribution profile variability among all the measurements (bottom graph). A p-value <0.01 in one region indicates there is a significant difference and a p-value >0.05 indicates no significant difference meaning ether that the compared profiles are highly the same in the 2 groups or the compared profiles are highly variable in the 2 groups.

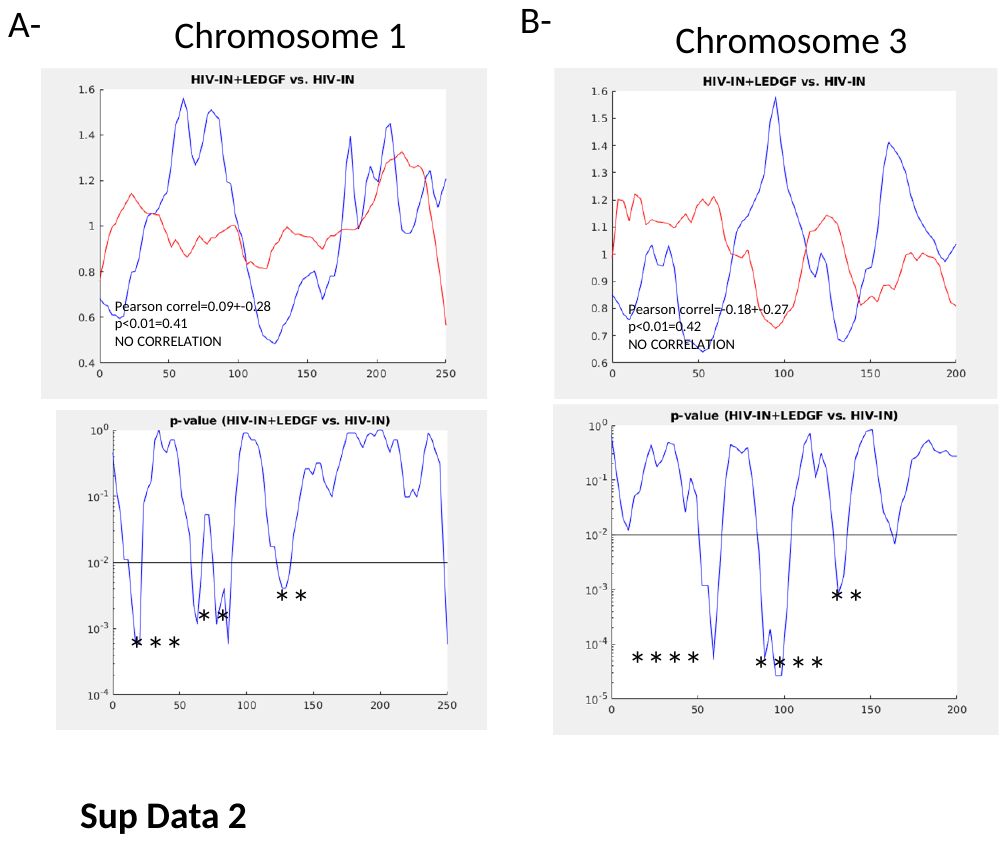

**S4. Functional association of WT, D253H and R231H IN CTD mutants to mononucleosomes.** Recombinant purified WT, mutant HIV-1 INs (10pmoles of IN monomers) were incubated MNs (250ng, *i.e.,* 125ng DNA), in 10µl interaction buffer (50mM HEPES, pH7.5; 1µg/ml BSA;1mM DTT;0.1% Tween 20;10% glycerol;and 50 to 240mM NaCl) for 20min on ice and then for 30min at room temperature. A 12.5µl aliquot of DynabeadsMyOne Streptavidin T1 (Invitrogen, ref. 65601) was then added to a total volume of 300µl interaction buffer and incubated at room temperature for 1h under rotation. The beads were washed three times with 300µl interaction buffer, and the precipitated products were resuspended in 10µl of Laemmli buffer, after which they were separated on a 12% gel via SDS-PAGE. Bound IN was detected by western blotting using a polyclonal anti-IN antibody and quantified as reported in (**A**) as the percentage of input precipitated under each condition. Integration assays were performed on MN (50ng in DNA) immobilized on streptavidin beads with 400nM of WT or mutated INs and 10nM of 17 bp of a 5’-radiolabeled viral U5 end. The percentage of integrated product was measured as indicated in materials and methods section and is shown in (**B**). All values are shown as the mean ± standard deviation (error bars) of three independent sets of experiments.

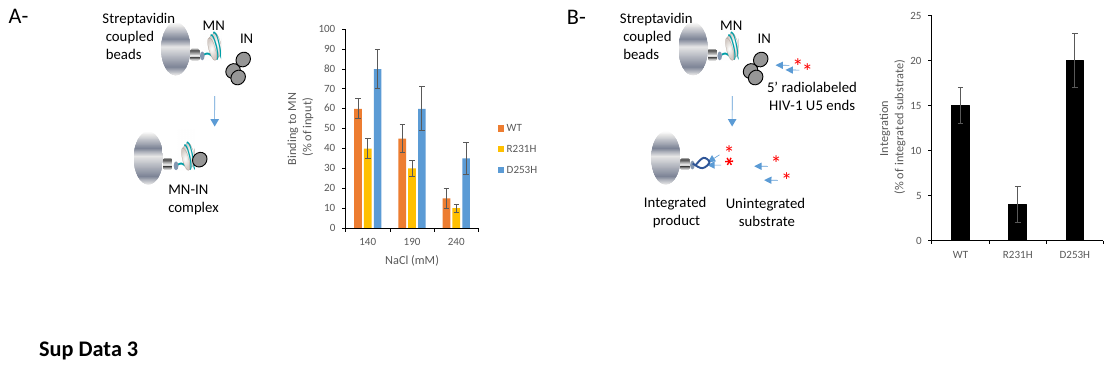

**S5. Assembly, purification and biochemical characterization of MVV and PFV intasomes.** FITC coupled intasomes were assembled and purified by size exclusion chromatography, (**A**) for MVV and (**C**) for PFV, following previously reported conditions (see materials and methods section). Integration assays were performed on naked p481 plasmid (Benleulmi et al. 2015) (**B** for MVV and **E** for PFV). Integration products were detected on 0.8% agarose gel. Integration product structures are reported as full site integration (FS product). o.c. open circular, s.c. supercoiled.

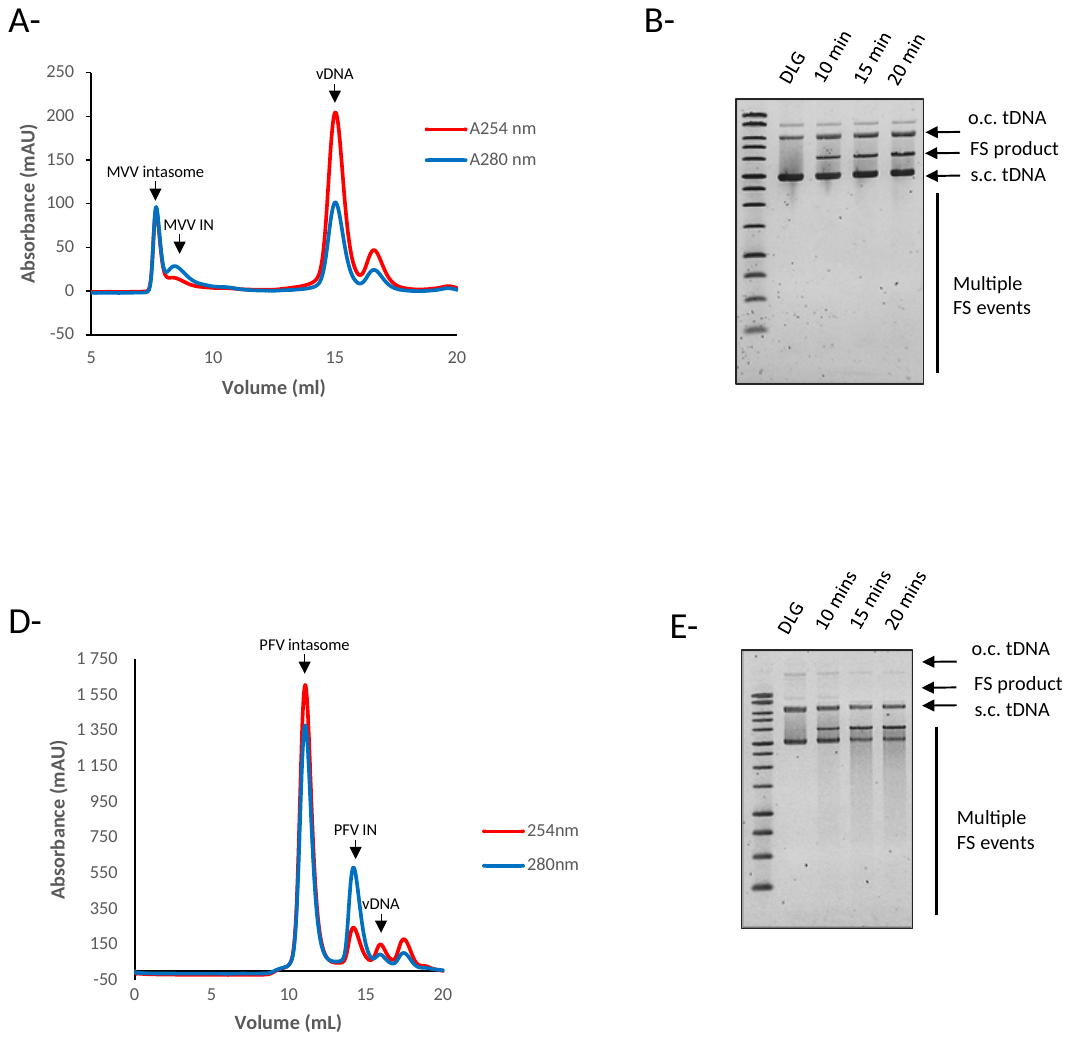

**S6. Analysis of the MVV and PFV intasome binding to chromosome 1.** Pearson correlation was computed for the MVV and PFV distribution curves and the proportion of significant correlation (p value) was provided >0 if correlation, =0 if no correlation (top graph **A**). The mean correlation +/- standard deviation is provided among all the comparison and a positive mean correlation indicates that most of the time, the curve correlates together, a close to 0 mean can indicate that curves do not correlate together. The p-value was also calculated as ranksum tests at each position along the chromosome in order to identify regions where there is a significant difference while taking into account the distribution profile variability among all the measurements (bottom graph **A**). A p-value <0.01 in one region indicates there is a significant difference and a p-value >0.05 indicates no significant difference meaning either that the compared profiles are highly the same in the 2 groups or the compared profiles are highly variable in the 2 groups. The Chromosome 1 distribution profile or PFV intasome in the presence of LEDGF/p75 was compared to MVV intasome distribution in **B**. The data are reported as means from the quantification of eight to nine chromosomes ± standard deviation (SD).

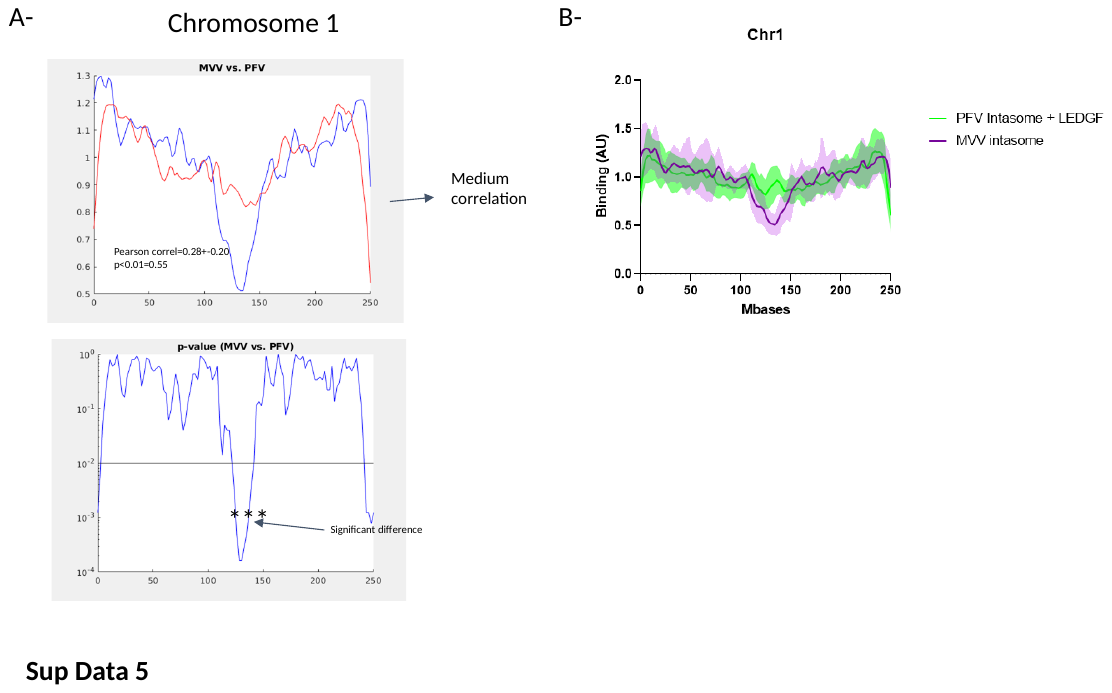

**S7. Sequence of the U5 ODN used for intasome assembly.** Rev and FITC-coupled fwd ODN were hybridized before intasome assembly by 85°C heating for 3min and slow decrease at room temperature overnight. The corresponding INs were then mixed with the ODN under conditions reported in the materials and methods section and intasome was purified by size exclusion chromatography.

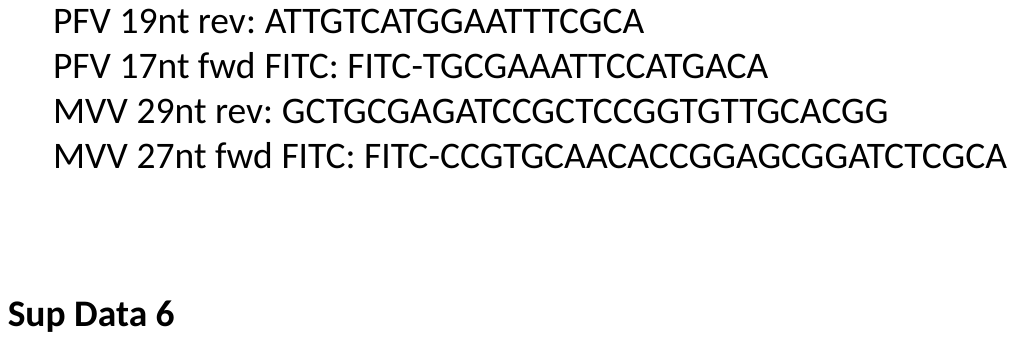
